# Biographika: rich interactive data visualizations on the web for the research community

**DOI:** 10.1101/021063

**Authors:** Pablo Pareja-Tobes, Eduardo Pareja-Tobes, Marina Manrique, Eduardo Pareja, Raquel Tobes

**Affiliations:** oh no sequences! research group, Era7 bioinformatics

## Abstract

When visualizing scientific data one of the current bottlenecks is the lack of interactivity. There already exist many options to build static data visualizations such as R, Matlab or Microsoft Excel among others. On the other hand, we can also find many different pieces of software with a broader or more specific aim that however must be installed locally. There is therefore a gap that is not covered by any of these latter two worlds. Here is where Biographika comes in handy since it provides scientists in general, and more specifically bioinformaticians, with a way of being able to use interactive rich visualizations on the web as part of their daily research. This first version of Biographika includes a set of charts combining diverse approaches that are thought to give users different perspectives of their research data. For the sake of interoperability and expressivity among other reasons we are using D3.js [1], the de facto standard visualization JavaScript library for manipulating documents based on data. But not only that, we incorporate new approaches as the fact of having fully interactive 3D charts that can be easily integrated with the rest of visualizations; providing the possibility of analyzing multidimensional data in a way that could otherwise be difficult to tackle. For that we use X3DOM[2], an open-source framework and runtime for 3D graphics on the Web that eases the integration of HTML5 and declarative 3D content. And last but not least, Biographika is also conceived as an effort to provide the data visualization layer for Bio4j [3] that researchers have been lacking in the past few years.

## Introduction

Many different options can be found for the visualization of data on the web. However, once we distinguish between static and interactive visualizations, the amount of options decreases drastically to just a few in the case we go for the latter. In addition to this, when talking about biological data, visualization tools or examples seem to be way behind compared to other fields such as for instance analytics. This is nevertheless something that should be fixed the sooner the better since the complexity of biological data is very high and, in some cases, being able to explore it by means of an interactive visualization could help to open the door to new scientific breakthroughs. Moreover, interactive data visualizations can in fact be linked together such that interactions in one visualization cause updates in another visualization. This technique is referred to as “multiple linked views” [4] and “brushing and linking” [5], [6]. This technique overcomes limitations of single visualizations by supporting exploration of the data through interaction. More information can be presented to the user with multiple linked views as compared to static visualizations. [7]

This, hence, was a strong motivation for us to create Biographika along the same lines as Bio4j, being both an effort to bring innovative cutting edge technologies to a field that unfortunately is lagging behind in this sense. It is important to state that Biographika was thus not thought as an isolated general tool for visualizing biological data but mainly as a visualization layer for data extracted from Bio4j.

Bio4j provides in fact a platform for the integration of semantically rich biological data using typed graph models, modeling and integrating most publicly available data linked with proteins into a set of interdependent graphs. These data include:

- UniProt [8]
- Gene Ontology [9]
- UniRef [10]
- Enzyme DB [11]
- NCBI Taxonomy [12]

Bio4j represents a unique resource for the current Bioinformatician, providing at once a solution for several key problems including: data integration; expressive, high performance data access; and a cost-effective scalable cloud deployment model. [3] The combination of these two projects therefore aims to fill an important gap in the world of bioinformatics data analysis.

## Our approach

Virtually all visualizations that are either developed or planned to be included in the near future as part of Biographika are based in D3.js to a moderate or greater extent. That way, rather than hiding the underlying scene-graph within a toolkit-specific abstraction, we are allowed to directly inspect and manipulate the standard DOM *(document object model)* when necessary. Users are then free to combine all of the data visualizations provided by Biographika with any other Javascript library whilst keeping full control over the graphical elements that compose the visualization. Once this has been said, it is also important to highlight how we use a different combination of d3.js based toolkits together with other more general Javascript libraries depending on the specific needs of each case. The list of libraries currently used include among others:

- JQuery [13] + JQuery UI [14]
- DC.js [15]
- Dimple.js [16]
- Crossfilter.js [17]
- X3DOM [2]
- Bootstrap [18]
- Slickgrid [19]
- Parallel coordinates [20]

Now we proceed to describe the various kinds of visualization utilizing the combination of strategies aforementioned.

### Dimple.js based charts

A set of simple but powerful charts were implemented using dimple.js, *“an object-oriented API for business analytics powered by D3”*[16]. One of the main features from this API is that it provides a gentle learning curve and minimal code to achieve something productive. Also when using dimple.js, d3 objects are still exposed in a way that they can be both picked up and run in order to create more complex and complete results.

We are now presenting the different charts that were implemented using this technology *(be aware that all of them include tooltips when hovering over the data showing the respective values. Also it is important to highlight that adding any extra functionality based on mouse interaction with the charts could be very easily integrated).*

#### Pie chart

In this example the pie chart is displaying the different percentages regarding the taxonomic assignment of a set of samples after some process has been carried out. This is a good way to see the big picture for the distribution of a set of groups over the total. (See Figure 6)

#### Stacked bar chart

Figure 1 indicates the abundance of species through a set of samples that would have been analyzed in a specific experiment. This visualization is very useful to see the inner distributions of species in specific samples whilst having the possibility of comparing them to the rest of samples included in the analysis.

**Figure 1:**
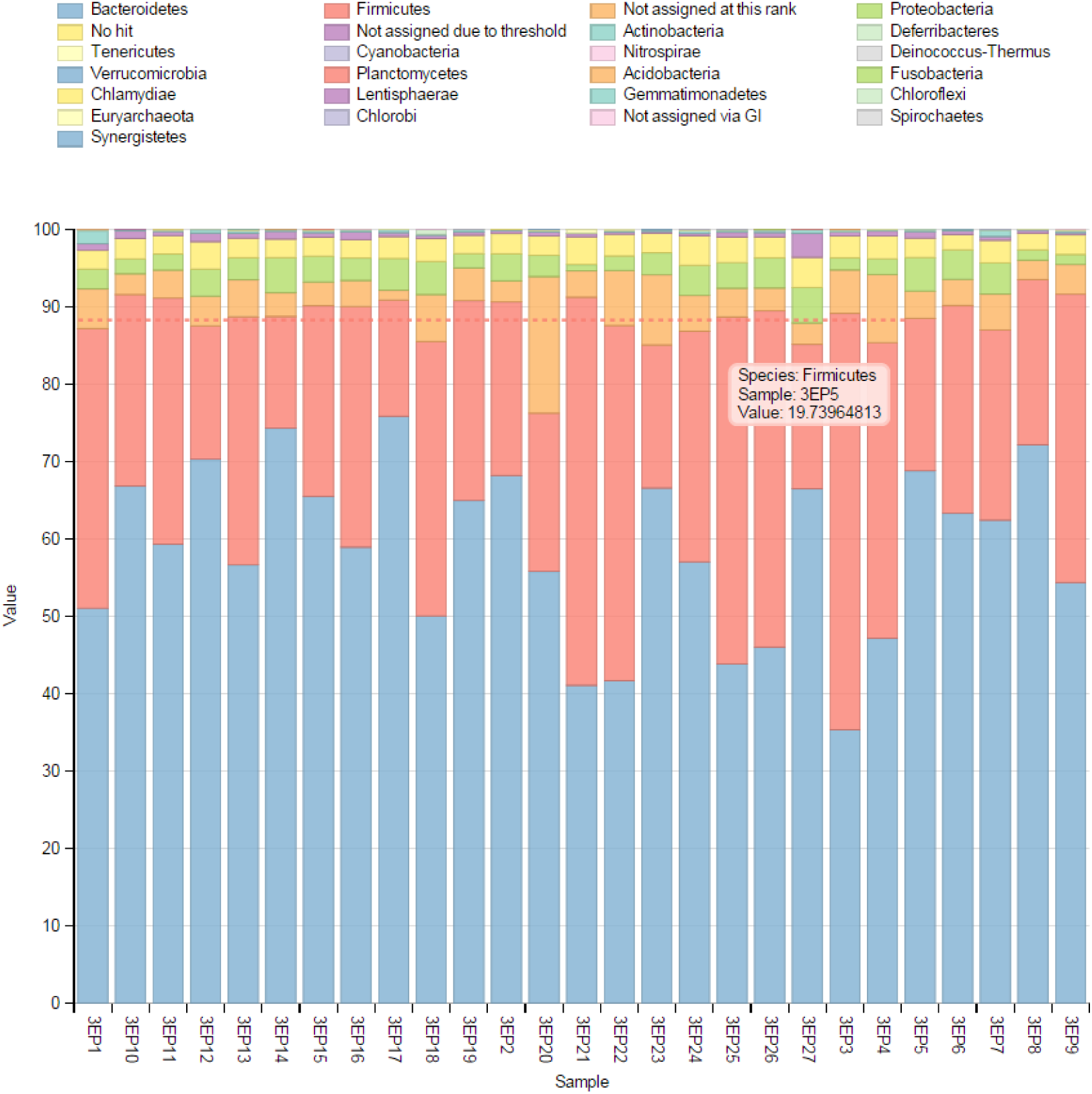
Representing the abundance of species through a set of samples by means of a stacked bar chart.

#### Multibar chart

In this case samples have been grouped by some criteria in four different groups. We are then getting insights on values for these groups over the most abundant species. This chart, whilst not being the same as the stacked bar chart, has a similar role in the sense that we can have both an inner and outer look at the data at the same time. (See Figure 2).

**Figure 2:**
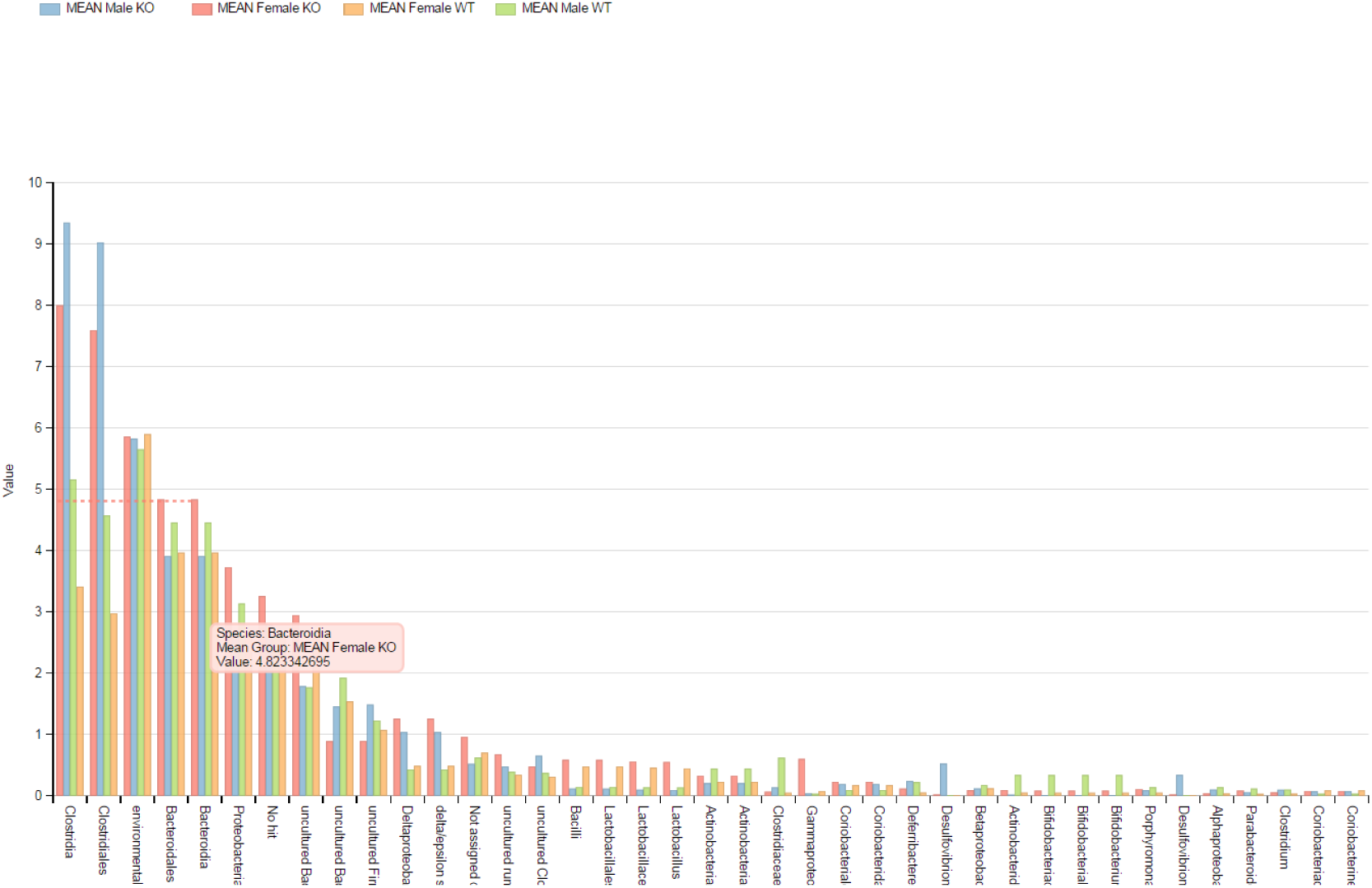
Multibar chart showing the abundance of species through a set of samples.

#### Multiline chart

The multiline chart could be seen as a variation of the example aforementioned since it is showing the same data as the multibar chart but from a different perspective. In this case sample groups are represented as lines and thus we have the possibility of having a much clearer idea of the fluctuations in value over the different species represented. (See Figure 3)

**Figure 3:**
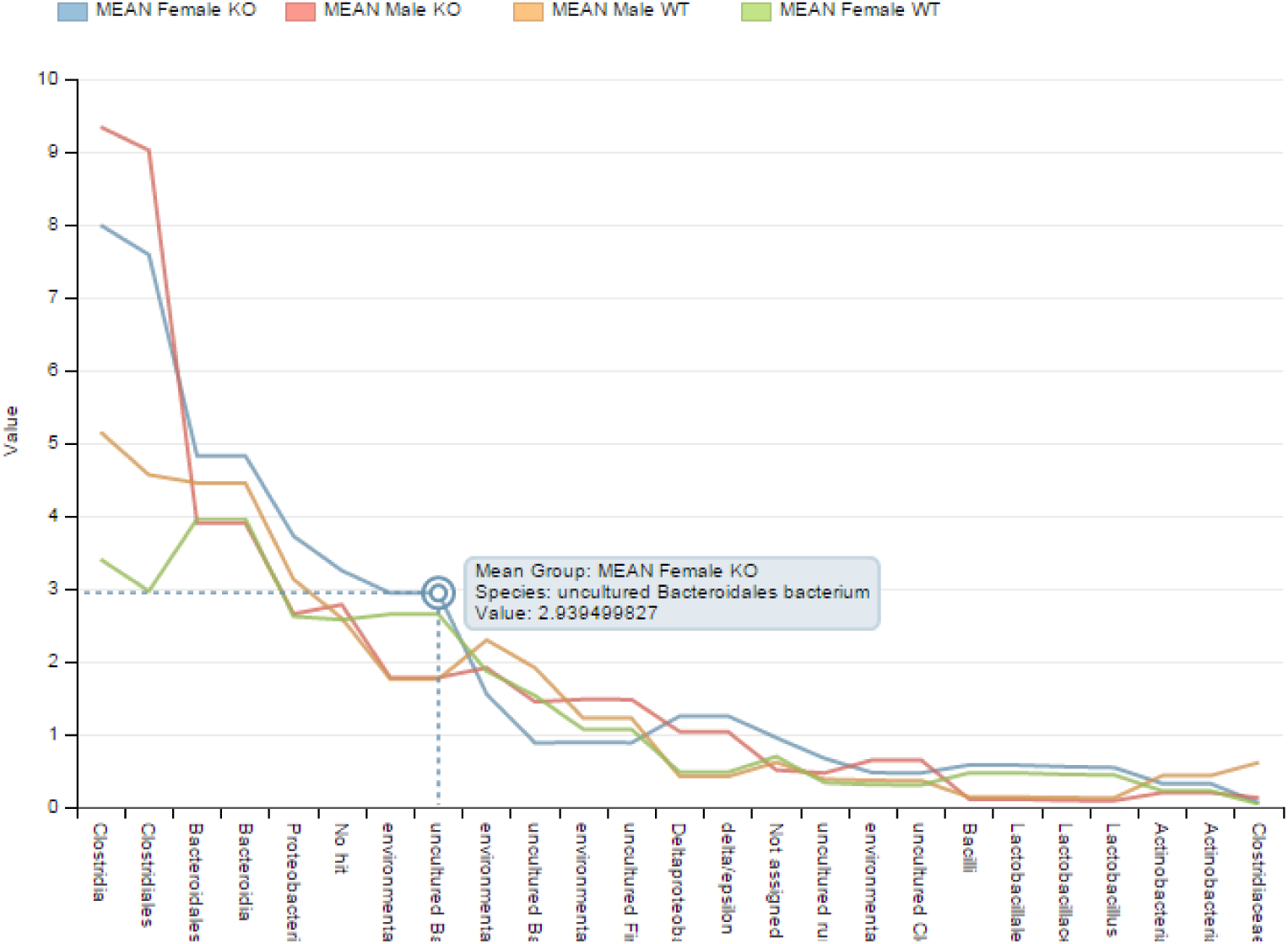
Multiline chart showing the fluctuations in value over the set of species represented.

### Parallel coordinates chart

This data visualization is built using sytagmatic’s Parallel Coordinates (0.6.0) *(A d3-based parallel coordinates plot in canvas)* [20] plus slickgrid *(A lightning fast JavaScript grid/spreadsheet)* [19]

According to Wikipedia:

*Parallel coordinates is a common way of visualizing high-dimensional geometry and analyzing multivariate data. To show a set of points in an n-dimensional space, a backdrop is drawn consisting of n parallel lines, typically vertical and equally spaced. A point in n-dimensional space is represented as a polyline with vertices on the parallel axes; the position of the vertex on the i-th axis corresponds to the i-th coordinate of the point. This visualization is closely related to time series visualization, except that it is applied to data where the axes do not correspond to points in time, and therefore do not have a natural order. Therefore, different axis arrangements may be of interest.*

In this specific example parallel coordinates were utilized for the visualization and comparison of panproteome sequences. We implemented a customizable color function that gave us at first sight an idea of how much were panproteome proteins shared along the different panproteomes. Having the possibility to rearrange the ordering of the axes was also key in some cases to better narrow down and compare specific proteins we wanted to focus on. Finally we should mention how, even though it was used for panproteome analysis in this case, this visualization could be adapted for any other use case extremely easily.

Any multivariable CSV file could in principle be visualized with our implementation, only the following lines of Javascript code should be modified in order to change the specific columns of data to be displayed as axes:

~~~
**var** dimensions = [
      ′column1′,
      ′column2′,
      ′column3′
];
**var** types = {
      ″column1″ : ″number″,
      ″column2″ : ″number″,
      ″column3″ : ″string″
};
~~~

The array dimensions defines the names of the columns to be included. On the other side types simply indicates the data type for each of the columns previously defined.

In additon to this, it is important to highlight the features that make this tool **highly interactive**. A selection or filtering can be performed both on the rows of the table and the parallel coordinates chart, having any of the two previously mentioned updated in real time depending on the selection performed on the other. Besides we included a couple of buttons that can switch the background of the visualization to better appreciate specific line connectors on different cases. (See Figure 4)

**Figure 4:**
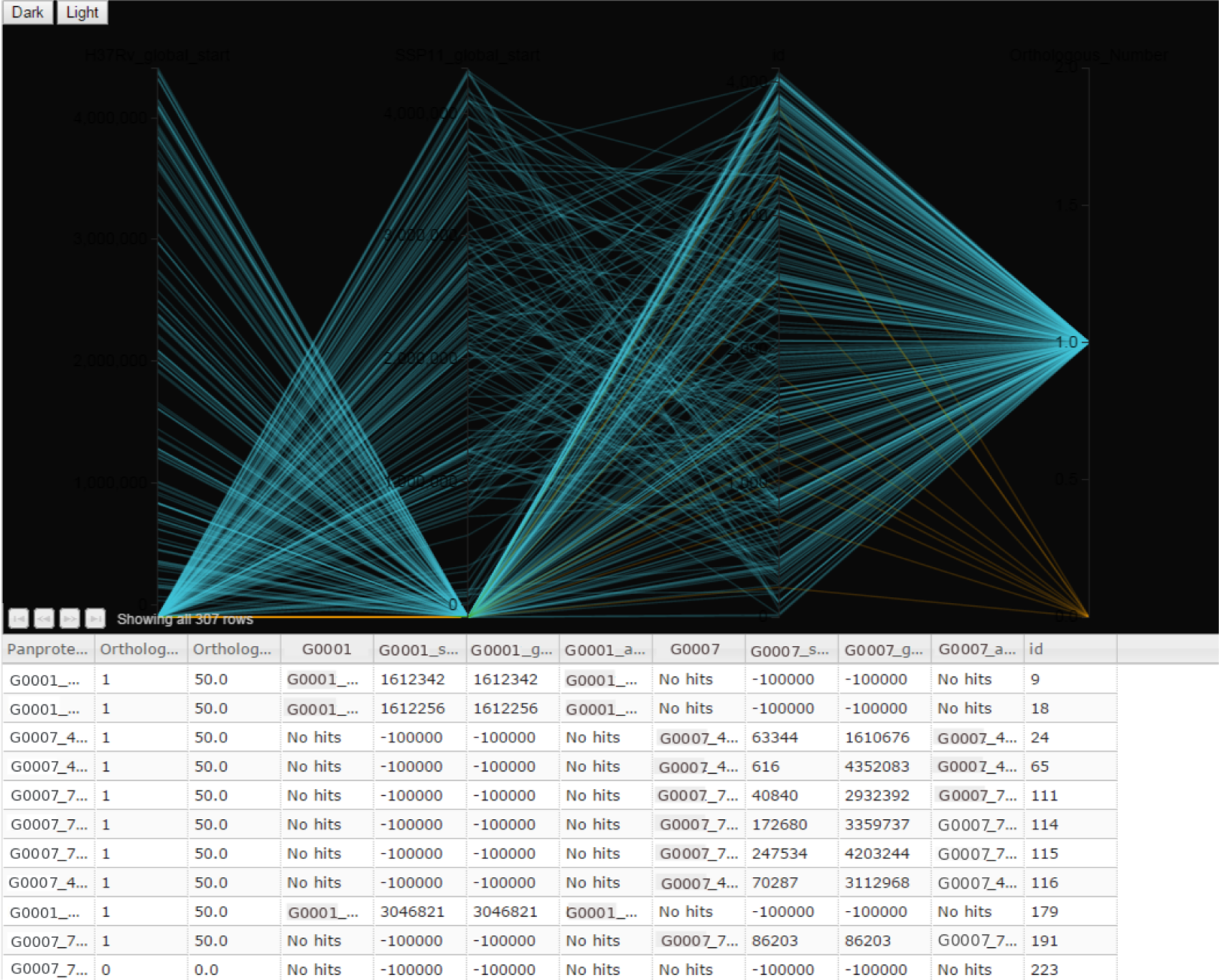
Parallel coordinates visualization.

### Combined approach: taxonomy tree plus bar chart

We also explored combined approaches that have been proved to be a great fit in cases where simpler diagrams could not cope with the complexity of the data that was to be visualized. Here we are putting together a variation of the collapsible tree d3.js block from Robert Schmuecker with a dimple.js bar chart and JQuery UI — *a curated set of user interface interactions, effects, widgets, and themes built on top of the jQuery JavaScript Library.*[14]

This chart includes a zoomable, collapsible taxonomy tree with auto-sizing that holds all the taxa involved in the analysis. Branches can be either collapsed or opened by clicking on the taxa nodes. Meanwhile when clicking on the name of a specific taxon a resizable dialog will be opened displaying a bar chart representing the percentage of assignments to this taxon for all the samples that were included in the analysis. (See Figure 5)

**Figure 5:**
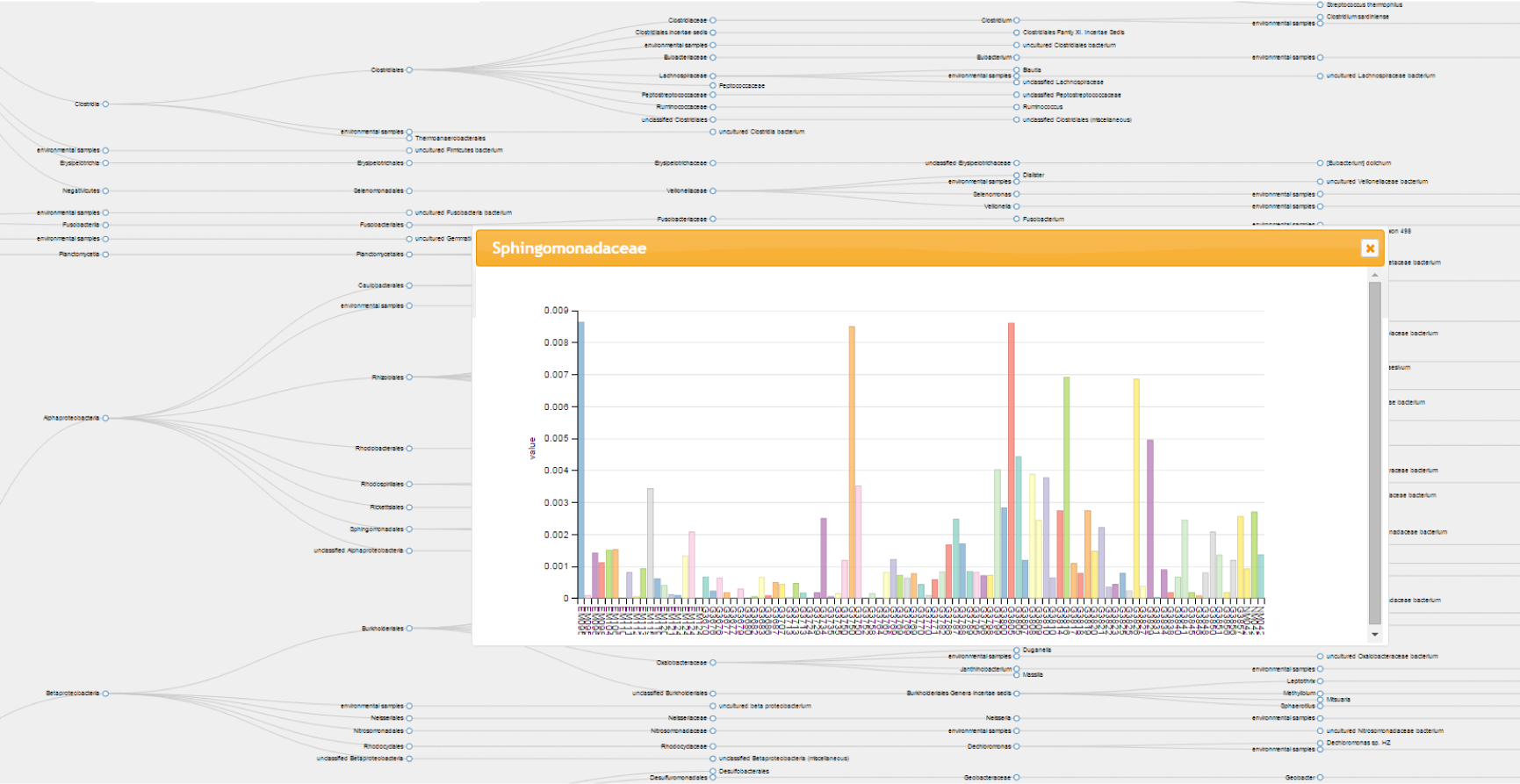
An example of a combined approach: an interactive zoomable taxonomic tree plus a bar chart included in a dialog shown by clicking on taxa nodes.

**Figure 6:**
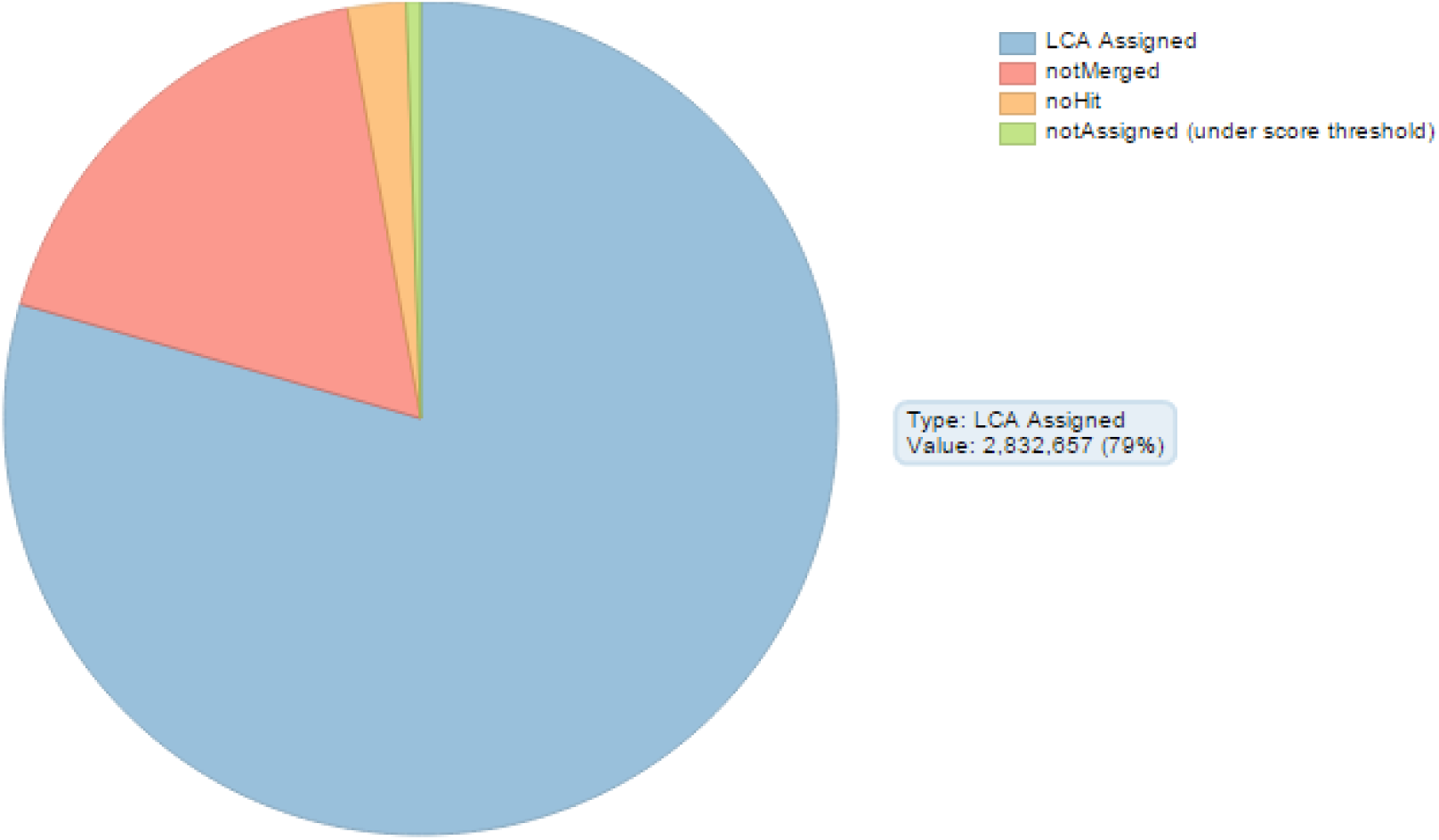
Pie chart showing the different percentages of a set of groups over the total.

### Customizable multicolor heat map

We also provide a heat map visualization including a special feature which is a customizable multicolor function. This color function consists of three different intervals [0,a),[a,b),[b,100] where both a and b values can be customized by the user having the heat map updated in real time according to these new values. Having the possibility to define these three intervals allows the user to explore the abundance at three different scales, which in turn makes possible to visualize representative values that may not be the highest of the heat map. (See Figure 7)

**Figure 7:**
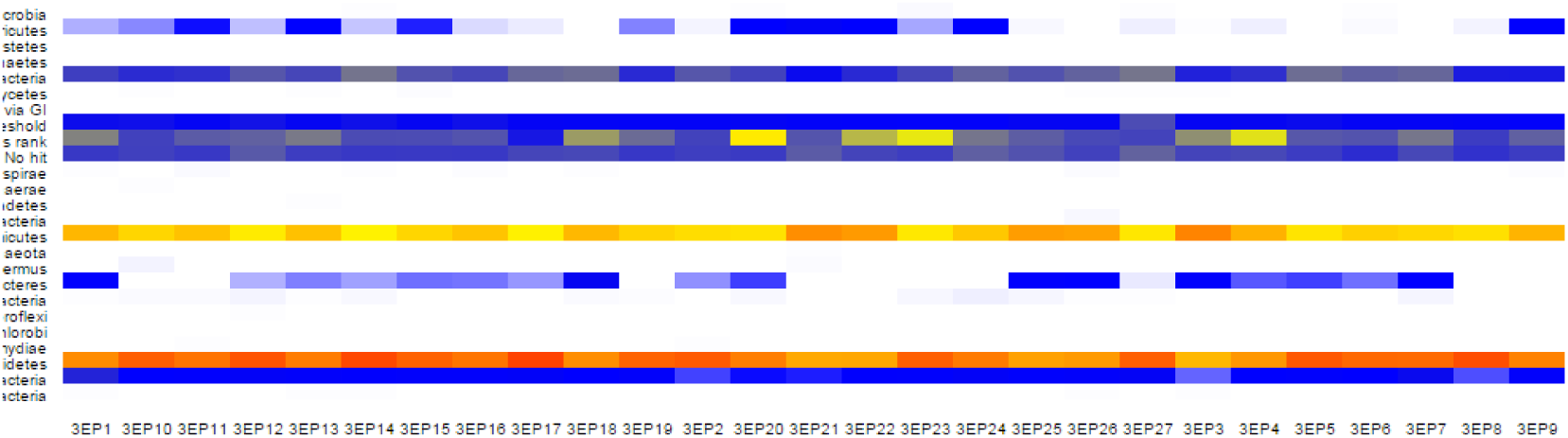
Heat map including a customizable color function based in three different intervals defined by the user.

### Antibiotic Resistance Explorer

This is another example of an integrative combined approach with the aim of having a dashboard where scientists can explore a set of variables around the concept of antibiotic resistance for a set of genes associated to a taxon and its descendants. In order to build this tool we are using the following technologies:

- **DC.js**: *a javascript charting library with native crossfilter support and allowing highly efficient exploration on large multi-dimensional dataset. It leverages d3 engine to render charts in css friendly svg format. Charts rendered using dc.js are naturally data driven and reactive therefore providing instant feedback on user’s interaction.* [15]
- **Crossfilter.js** *a JavaScript library for exploring large multivariate datasets in the browser. Crossfilter supports extremely fast (<30ms) interaction with coordinated views, even with datasets containing a million or more records.* [17]
- **Bootstrap** *an open source HTML, CSS, and Javascript framework for developing responsive, mobile first projects on the web. By using this technology we can easily and efficiently scale our tools and data visualizations while using a single code base for all devices.* [18]

The main feature of this tool is that the user can filter by clicking in any of the charts and every other chart will be re-rendered dinamycally according to the filtering specified on the chart that was clicked. There also exist a set of *‘reset’* buttons *(one for each chart)* that allow to clear the filterings that may have previously been applied to the respective chart.

As it can be seen in the Figure 8 the visualization includes four different charts: a pie chart and three bar charts. The pie chart represents the proportion of taxa that, depending on the percentage of proteins associated which are annotated as antibiotic resistance, are classified as:

**Figure 8:**
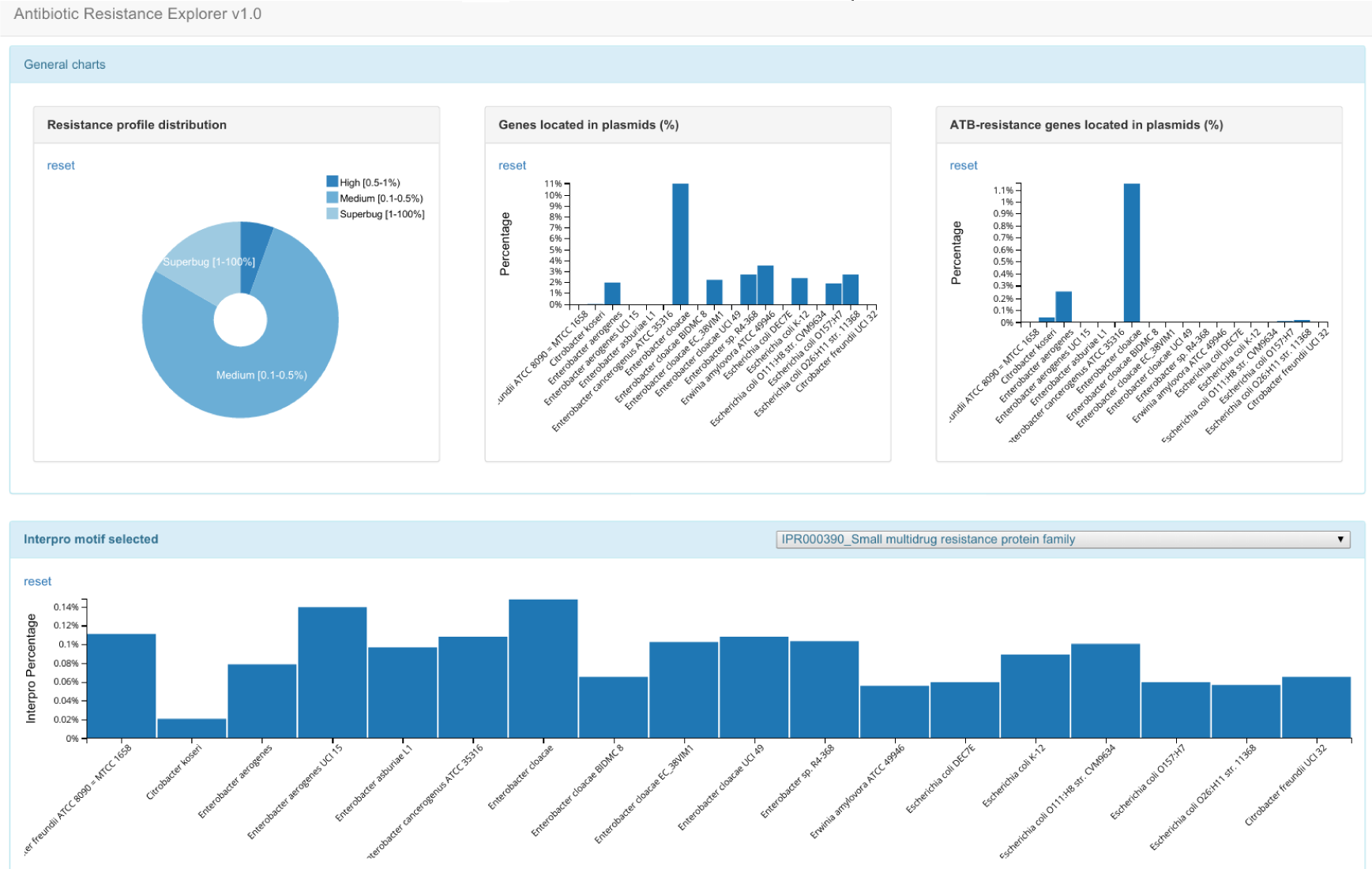
Antibiotic Resistance Explorer.

- **Low** [0 - 0.1%)
- **Medium** [0.1% - 0.5%)
- **High** [0.5% - 1%)
- **Superbug** [1%-100%]

Together with the pie chart, two bar charts are included in the same panel. The first one shows the percentage of plasmid proteins found per taxa whereas the second shows the percentage of proteins annotated both as plasmid and antibiotic resistance.

The panel below includes at the right end a drop-down list that is dynamycally populated with as many Interpro motifs as columns are found for them in the CSV file used as input for this data visualization. Thus, by using this element we can visualize the percentage of proteins per taxa that are associated to antibiotic resistance related Interpro motif selected by the user. Interacting with this chart will also filter the data that is being visualized, updating the rest of charts accordingly to the selection.

### 3D Bar Chart

Finally we have included a very useful visualization for multivariate data by representing a 3 dimensional matrix of values with species and samples as axes. As it was mentioned in the abstract, in order to build this chart we use X3DOM[2] for the 3D visualization. The bar chart is inspired by camio’s D3.js block, although it has been largely improved in order to have an interactive 3D matrix bar chart instead. See Figure 9.

**Figure 9:**
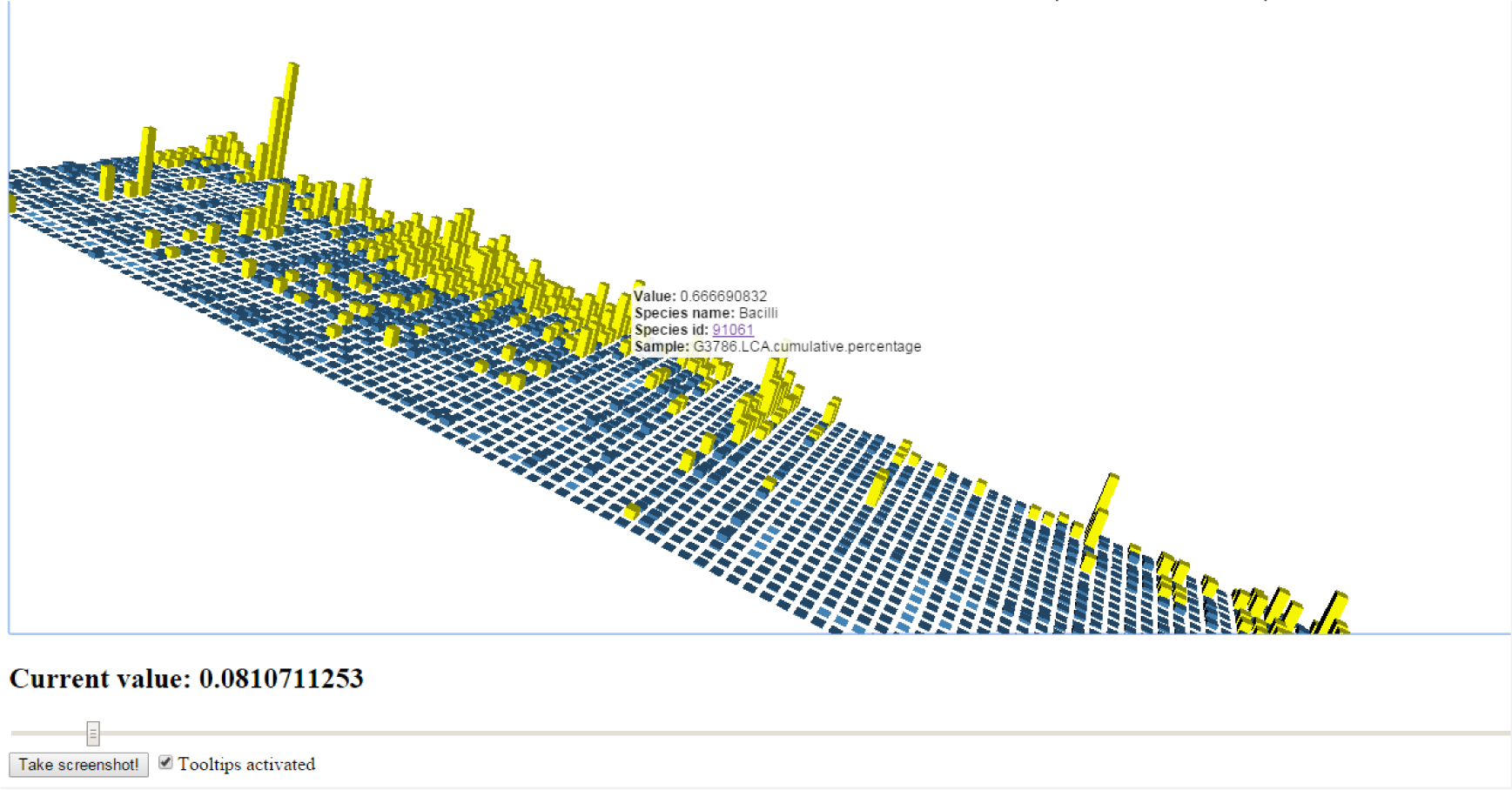
Bar Chart representing a 3 dimensional matrix of values with species and samples as axes.

This chart includes several features that allow the user to see different characteristics of the data by interacting with the visualization. First of all we have added tooltips to the bars that can be either activated or deactivated by means of a checkbox. These tooltips include data attributes such as: sample name, species name, percentage value, and a clickable piece of text holding the species id that opens a new tab leading to the NCBI taxonomy official page for that specific taxon.

Apart from that there also exists a slider by which the user is able to highlight all bars that hold a value greater than that specified by the slider. This is a good way to quickly get a general idea of the distribution of pairs species-sample that have values above and below the desired threshold.

Finally we also added a button that takes a screenshot of the current state of the chart allowing researchers to easily have an image showing their results that could later be used for publication or other purposes.

## Biographika codebase

Biographika is open source, licensed under the open source approved AGPLv3 license. All development happens on GitHub, under the Biographika organization.

## References

[1] M. Bostock, V. Ogievetsky, and J. Heer D^3^ data-driven documents Visualization and Computer Graphics, IEEE Transactions on 17 (12): 2301–2309. 2011.

[2] J. Behr, P. Eschler, Y. Jung, and M. Zöllner X3dom: A dom-based html5/x3d integration model In Proceedings of the 14th International Conference on 3D Web Technology ser. Web3D ’09 Darmstadt, Germany: ACM, 2009, 127–135. DOI: 10.1145/1559764.1559784.

[3] P. Pareja-Tobes, R. Tobes, M. Manrique, et al. Bio4j: A high-performance cloud-enabled graph-based data platform BioRxiv, 016758. 2015. DOI: 10.1101/016758.

[4] J. Dykes, A. MacEachren, and M. Kraak Exploratory visualization with multiple linked views 2004.

[5] D. Keim et al. Information visualization and visual data mining Visualization and Computer Graphics, IEEE Transactions on 8 (1): 1–8. 2002.

[6] L. Anselin, I. Syabri, and O. Smirnov Visualizing multivariate spatial correlation with dynamically linked windows Urbana 51, 61801. 2002.

[7] C. Kelleher and H. Levkowitz Reactive data visualizations In IS&T/SPIE Electronic Imaging International Society for Optics and Photonics 2015, 93970N–93970N.

[8] U. Consortium et al. Uniprot: A hub for protein information Nucleic acids research, gku989. 2014. DOI: 10.1093/nar/gku989.

[9] M. Ashburner, C. A. Ball, J. A. Blake, et al. Gene ontology: Tool for the unification of biology Nature genetics 25 (1): 25–29. 2000. DOI: 10.1038/75556.

[10] B. E. Suzek, H. Huang, P. McGarvey, et al. Uniref: Comprehensive and non-redundant uniprot reference clusters Bioinformatics 23 (10): 1282–1288. 2007. DOI: 10.1093/bioinformatics/btm098.

[11] A. Bairoch The enzyme database in 2000 Nucleic acids research 28 (1): 304–305. 2000. DOI: 10.1093/nar/28.1.304.

[12] E. W. Sayers, T. Barrett, D. A. Benson, et al. Database resources of the national center for biotechnology information Nucleic acids research 39 (Suppl 1): D38–D51. 2011. DOI: 10.1093/nar/gkq1172.

[13] Jquery URL: https://jquery.com/.

[14] Jquery ui URL: https://jqueryui.com/.

[15] Dc.js – dimensional charting javascript library URL: https://dc-js.github.io/dc.js/.

[16] Dimple.js – a simple charting api for d3 data visualisations URL: http://dimplejs.org/.

[17] Crossfiler.js – fast multidimensional filtering for coordinated views URL: http://square.github.io/crossfilter/.

[18] Bootstrap – the most popular html, css, and js framework for developing responsive, mobile first projects on the web. URL: http://getbootstrap.com.

[19] Slickgrid – a lightning fast javascript grid/spreadsheet URL: https://github.com/mleibman/SlickGrid/wiki.

[20] Parallel coordinates – a visual toolkit for multidimensional detectives URL: https://syntagmatic.github.io/parallel-coordinates/.

